# IL13Pred: A method for predicting immunoregulatory cytokine IL-13 inducing peptides for managing COVID-19 severity

**DOI:** 10.1101/2021.09.19.460950

**Authors:** Shipra Jain, Anjali Dhall, Sumeet Patiyal, Gajendra P. S. Raghava

## Abstract

Interleukin 13 (IL-13) is an immunoregulatory cytokine that is primarily released by activated T-helper 2 cells. It induces the pathogenesis of many allergic diseases, such as airway hyperresponsiveness, glycoprotein hypersecretion and goblet cell hyperplasia. IL-13 also inhibits tumor immunosurveillance, which leads to carcinogenesis. In recent studies, elevated IL-13 serum levels have been shown in severe COVID-19 patients. Thus it is important to predict IL-13 inducing peptides or regions in a protein for designing safe protein therapeutics particularly immunotherapeutic. This paper describes a method developed for predicting, designing and scanning IL-13 inducing peptides. The dataset used in this study contain experimentally validated 313 IL-13 inducing peptides and 2908 non-inducing homo-sapiens peptides extracted from the immune epitope database (IEDB). We have extracted 95 key features using SVC-L1 technique from the originally generated 9165 features using Pfeature. Further, these key features were ranked based on their prediction ability, and top 10 features were used for building machine learning prediction models. In this study, we have deployed various machine learning techniques to develop models for predicting IL-13 inducing peptides. These models were trained, test and evaluated using five-fold cross-validation techniques; best model were evaluated on independent dataset. Our best model based on XGBoost achieves a maximum AUC of 0.83 and 0.80 on the training and independent dataset, respectively. Our analysis indicate that certain SARS-COV2 variants are more prone to induce IL-13 in COVID-19 patients. A standalone package as well as a web server named ‘IL-13Pred’ has been developed for predicting IL-13 inducing peptides (https://webs.iiitd.edu.in/raghava/il13pred/).

**Key Points:**

- Interleukin-13, an immunoregulatory cytokine plays an important role in increasing severity of COVID-19 and other diseases.
- IL-13Pred is a highly accurate in-silico method developed for predicting the IL-13 inducing peptides/ epitopes.
- IL-13 inducing peptides are reported in various SARS-CoV2 strains/variants proteins.
- This method can be used to detect IL-13 inducing peptides in vaccine candidates.
- User friendly web server and standalone software is freely available for IL-13Pred

**Author’s Biography:**

1. Shipra Jain is currently working as Ph.D. in Computational Biology from Department of Computational Biology, Indraprastha Institute of Information Technology, New Delhi, India.
2. Anjali Dhall is currently working as Ph.D. in Computational Biology from Department of Computational Biology, Indraprastha Institute of Information Technology, New Delhi, India.
3. Sumeet Patiyal is currently working as Ph.D. in Computational Biology from Department of Computational Biology, Indraprastha Institute of Information Technology, New Delhi, India.
4. Gajendra P. S. Raghava is currently working as Professor and Head of Department of Computational Biology, Indraprastha Institute of Information Technology, New Delhi, India.

## Introduction

IL-13 is an immune-regulatory cytokine primarily secreted by activated T helper-Type (Th) 2 cells that inhibits inflammatory cytokine production [1,2]. In literature many studies have shown that IL-13 is also produced by diverse cell types, including eosinophils, mast cells, basophils, smooth muscle cells, natural killer cells, and fibroblasts with varied biological functions [3,4]. The transcription of IL-13 cytokine is mainly regulated by GATA3 transcription factor. IL-13 has approx. 25% sequence homology with IL-4 and is located on human chromosome 5q31 [4]. It has been shown that IL-4 and IL-13 are functionally related, but it is surprising that IL-13 seems to be a more promising target for designing therapeutics than IL-4 [3]. As shown in Figure 1, it mediates several vital functions in diverse biological pathways including regulation of airway hyperresponsiveness, allergic inflammation, mastocytosis, goblet cell hyperplasia, tissue eosinophilia, IgE Ab production, tumor cell growth, tissue remodeling, intracellular parasitism, and fibrosis.

**Figure 1:**
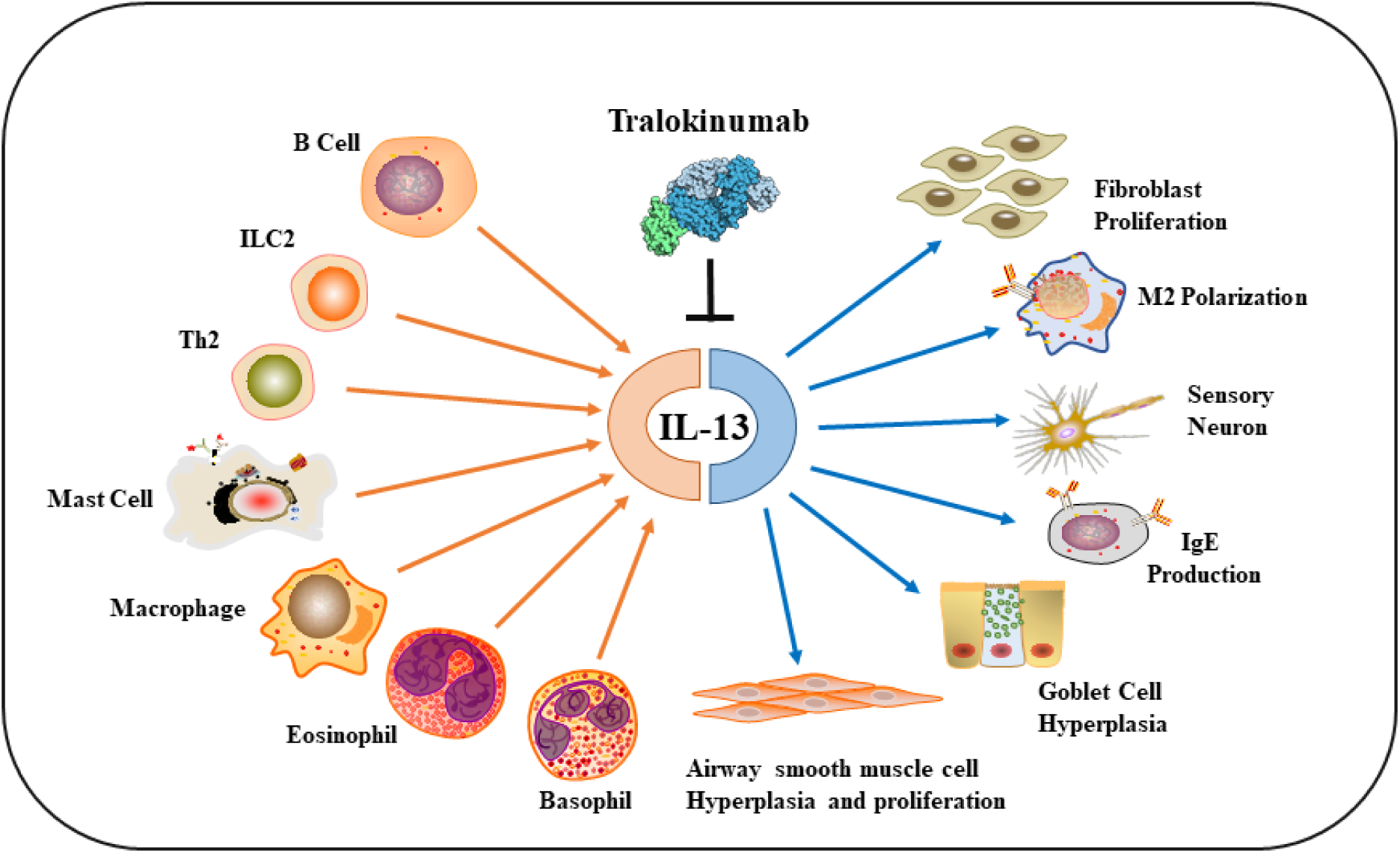
Schematic representation of IL-13 activators and its functions in biological pathways.

Many studies exhibited that alterations in IL-13 effector functions can be targeted to treat certain cancers, like B-cell chronic lymphocytic leukemia and Hodgkin’s disease [5,6]. It inhibits tumor immunosurveillance in homo sapiens, therefore IL-13 inhibitors can act as effective cancer immunotherapeutics candidates by activating type-1 anti-cancer defense mechanisms [7,8]. Moreover, studies have reported that IL-13/ IL-4 receptors have a crucial role in prognosis of cancers such as pancreatic, gastric, and colon cancer [9–12]. They interact with tumor microenvironment by activating tumor-associated macrophages and myeloid-derived suppressor cells [13,14]. IL-13 receptors were found to be overexpressed in glioblastoma multiforme human samples in situ, whereas, normal specimen expressed few IL-13 binding receptor sites [15]. Moreover, increased IL-13 levels were observed in pulmonary artery hypertension (PAH) patients as compared to non-PAH controls [16]. Involvement of IL-13 receptor α1 in myocardial homeostasis embarking its role in cardiovascular diseases [17].

IL-13 has emerged as a central regulator in airway hyperresponsiveness, fibrosis, mucus hypersecretion and switching of B cell antibody production from IgM to IgE. Research highlights that the IL-13 pathway could be a promising target in the treatment of diverse asthma phenotypes [18,19]. It has been shown in the past that anti-IL-13 drugs plays a major role in controlling the ‘Th2 high’ asthmatic phenotype [20]. Therefore, anti-IL-13 drugs have become popular (anrukinzumab, lebrikizunab and tralokinumab) for controlling the severity of asthma [21]. IL-13 also has crucial consequences on non-hematopoietic cells, including endothelial cells, smooth muscle cells, fibroblasts, epithelial cells, and sensory neurons [22–25]. Additionally, several studies have shown that the plasma levels of IL-13 were significantly higher in COVID-19 patients [26–28]. A recent study has shown that the COVID-19 infected patients with elevated IL-13 serum levels required ventilation support as compared to others. Also, COVID-19 patients prescribed with IL-13 inhibiting drug (Dupilumab) have shown less dreadful symptoms [26]. In another study, researchers have reported differential expression levels of 14 cytokines including IL-13 in healthy control, moderate and severe COVID-19 patients, and they observed that the higher IL-13 expression level is directly proportional to the COVID-19 severity [28]. Therefore, it is the need of the hour to develop a prediction method devoted to classify IL13 inducing peptides. To the best of our knowledge, IL-13Pred is the first of its kind to predict the IL-13 inducing and non-inducing peptides from its amino acid sequence.

In this study, we present an *in-silico* method to classify the IL-13 inducing and non-inducing peptides/epitopes. We have used experimentally reported IL-13 inducing and non-inducing peptides of humans from IEDB. Using this dataset, we have applied various state-of-the-art machine learning classifiers and the model performance was evaluated on the independent dataset. We have delivered a webserver and standalone version of ‘IL-13Pred’ for scientific community usage.

## Material and Methods

### Dataset Preparation and Pre-processing

Initially, we extracted 343 IL-13 inducing experimentally validated peptides of humans from IEDB [29]. During pre-processing, we have removed the peptides with length less than 8 or more than 35 amino acids [30]. Also, to avoid any redundancy in the dataset, we have removed duplicate copies of the peptides (if any). Eventually, we left with a positive dataset of 313 IL-13 inducing peptides. One of the major challenges was to compile a negative dataset of experimentally validated non-inducing peptides. To overcome this issue, we have acquired the negative dataset from the recently published article IL6Pred [30], and after the pre-processing we obtained 2908 non-inducing peptides. Finally, we proceeded with a positive dataset of 313 IL-13 inducing and a negative dataset of 2908 non-inducing unique peptides.

### Feature Generation

In the present method, we have implemented Pfeature to compute various types of descriptors using sequence information of the peptides. It generates a wide range of descriptors in a fixed vector size of a given protein or peptide sequence [31]. We utilized composition-based module of Pfeature for our dataset, and generated a vector of 9165 features for each sequence. For our dataset we computed 15 types of composition-based features like, AAC, DPC, TPC, ATC and many more as given in Table 1.

**Table 1:** List of descriptors with brief description and number of feature; computed using Pfeature.

| Description of Descriptor | Number of features or length of vector |
| --- | --- |
| Amino Acid Composition (AAC) | 20 |
| Dipeptide Composition (DPC) | 400 |
| Tripeptide Composition (TPC) | 8000 |
| Atomic and Bond Composition (ABC) | 9 |
| Physicochemical Properties Composition (PCP) | 30 |
| Residue Repeat Information (RRI) | 20 |
| Distance Distribution of Residue (DDR) | 20 |
| Shannon-Entropy of Protein (SEP) | 1 |
| Shannon Entropy of Residues (SER) | 20 |
| Shannon Entropy of Physicochemical Property (SPC) | 25 |
| Conjoint Triad Calculation (CTC) | 343 |
| Composition enhanced Transition Distribution (CTD) | 189 |
| Pseudo Amino Acid Composition (PAAC) | 21 |
| Amphiphilic Pseudo Amino Acid Composition (PAAC) | 23 |
| Quasi-Sequence Order (QSO) | 42 |
| Sequence Order Coupling Number (SOC) | 2 |

### Ranking and Selection of features

In order to extract the crucial features from a larger pool of features generated using Pfeature, we utilized SVC-L1-based feature selection technique from Scikit-learn package. SVC-L1 based method implements the support vector classifier (SVC) with linear kernel, penalized with L1 regularization [32]. This technique identifies the important features from the high-dimensional feature set. SVC-L1 is a faster method when compared to other available techniques for feature selection. Using this method, we have listed 95 important features from the pool of 9165 features.

Post that, feature-selector tool was implemented for ranking of the obtained key 95 features based on their performance in classifying the IL-13 inducing and non-inducing peptides. Decision tree-based algorithm light gradient boosting machine is implemented in the feature-selector tool which calculates the rank of feature based on the number of times a feature is used to split the data across all trees [33]. This tool generates top-ranked features (Supplementary Table S1) that were used to build machine learning prediction models for IL-13 inducing peptides.

### Cross-validation and External Validation dataset

We have implemented standard protocols to build, test and evaluate our prediction models. In the present study, we have applied both 5-fold cross-validation as well as external validation technique. The complete dataset was divided in 80:20 proportion, where 80% (i.e. 250 positive and 2326 negative) of the dataset was categorized as training and the prevailing 20% (i.e. 63 positive and 582 negative) was designated as independent dataset [34].

### Machine learning Models and Evaluation

In order to develop a prediction method for classifying IL-13 inducing peptides, we have implemented diverse machine learning techniques. In this study, we have used various classifiers such as eXtreme Gradient Boosting (XGB), K-nearest neighbor (KNN), support vector machine (SVM), gaussian naive bayes (GNB), decision tree (DT), linear regression (LR), and random forest (RF). We implemented Scikit-learn package of Python to build these machine learning prediction models [35].

Standard norms of five-fold cross validation were followed such as marking four folds of data as training dataset and remaining one-fold as testing data. We have repeated this step five times and have reported the average scores of five cycles in result section. We have optimized various parameters respective to each classifier to obtain the best performing models. In result section we have reported the standard measures used to estimate the performance of the developed prediction models. We have calculated both threshold-dependent as well as threshold-independent parameters for evaluating the performance. Sensitivity, specificity, accuracy and Matthews correlation coefficient (MCC) are calculated as threshold-dependent parameters using equations 1-4. Area Under the Receiver Operating Characteristic curve (AUC) is calculated as the threshold-independent measure.

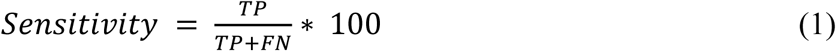

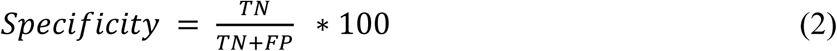

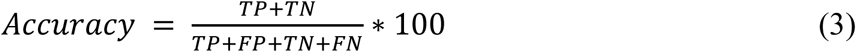

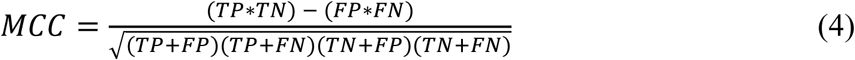

Where, FP is false positive, FN is false negative, TP is true positive and TN is true negative, respectively.

### Standalone and Web Server

We have provided a user-friendly web interface named ‘IL-13Pred’ to predict IL-13 inducing and non-inducing peptides (https://webs.iiitd.edu.in/raghava/il13pred). The web server is easy to use and has four major modules named as “Predict”, “Design”, “Protein Scan”, and “Blast Scan”. The front-end of the web interface was created using HTML5, JAVA, CSS3 and PHP scripts. This platform is compatible with most of modern devices including mobile, tablets, desktop and laptops. Standalone version of IL-13Pred can be downloaded from the provided URL https://webs.iiitd.edu.in/raghava/il13pred/stand.html.

## Results

### Compositional Analysis

We have computed the amino acid composition for the positive and negative dataset independently. The average amino acid composition for IL-13 inducing and non-inducing peptides is shown in Figure 3. The average composition of L, K, M and N residues are higher in IL-13 inducing peptides. Whereas, the amino acids G, P, Y and T residues are abundant in non-inducing peptides.

**Figure 2:**
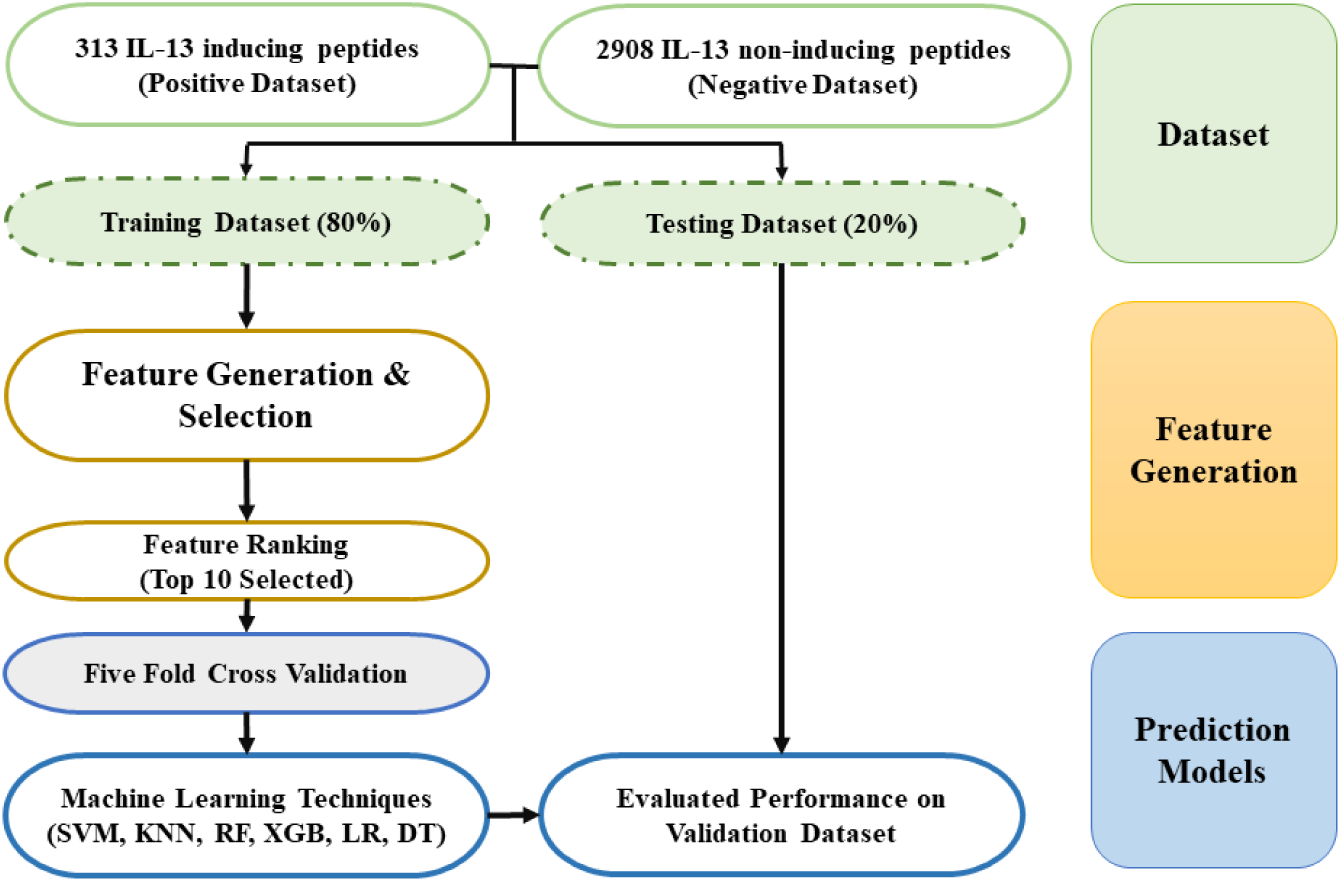
Work flow of IL-13Pred method algorithm.

**Figure 3:**
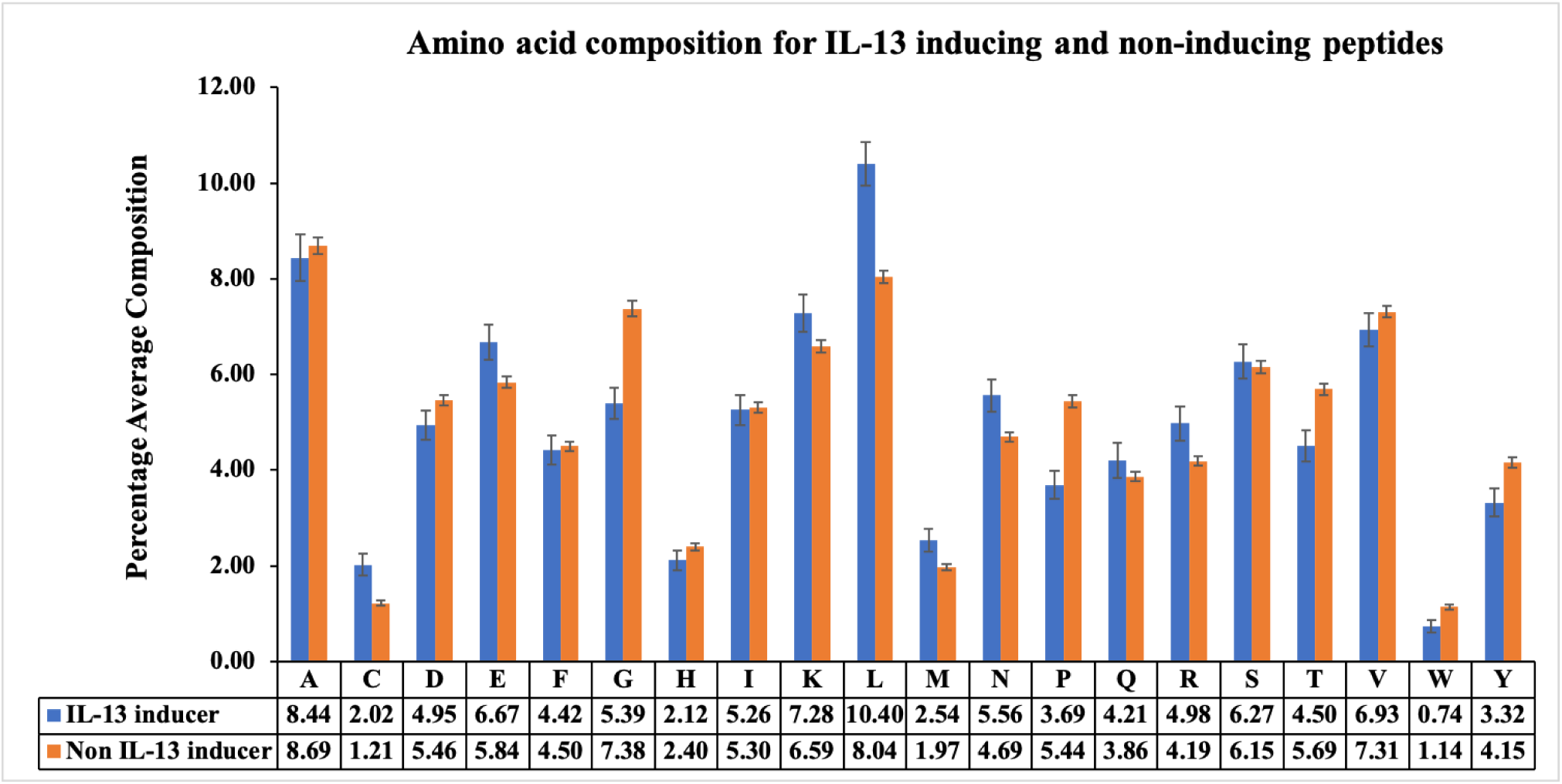
Average AAC of IL-13 inducing and non-inducing peptides.

### Positional Analysis

The amino acid positional analysis depicts the dominant residues at a particular position in IL-13 inducing and non-inducing peptides i.e., positive and negative dataset. In figure 4, two sample logo shows the important amino acid residue at a particular position and its relative occurrence in the sequence. The first eight positions represent the eight N-terminal peptide residues, and the last eight positions represent the eight C-terminal peptide residues. In IL-13 inducing peptides, we observe that Leucine (L) occurs more frequently at 2^nd^, 5^th^, 7^th^,9^th^, 11^th^ and 16^th^ position. Further, Asparagine (N) is mostly preferred at 6^th^, 8^th^, 14^th^ and 16^th^ position. Whereas, in negative dataset, we observe that the Threonine (T) occurred at 1^st^, 6^th^, 13^th^, 16^th^ position with good abundance, and Proline (P) preferred at 3^rd^, 5^th^, 14^th^ position. In addition to these two amino acids, Glycine (G) showed an interesting pattern in duplets at 7^th^ and 14^th^ position.

**Figure 4:**
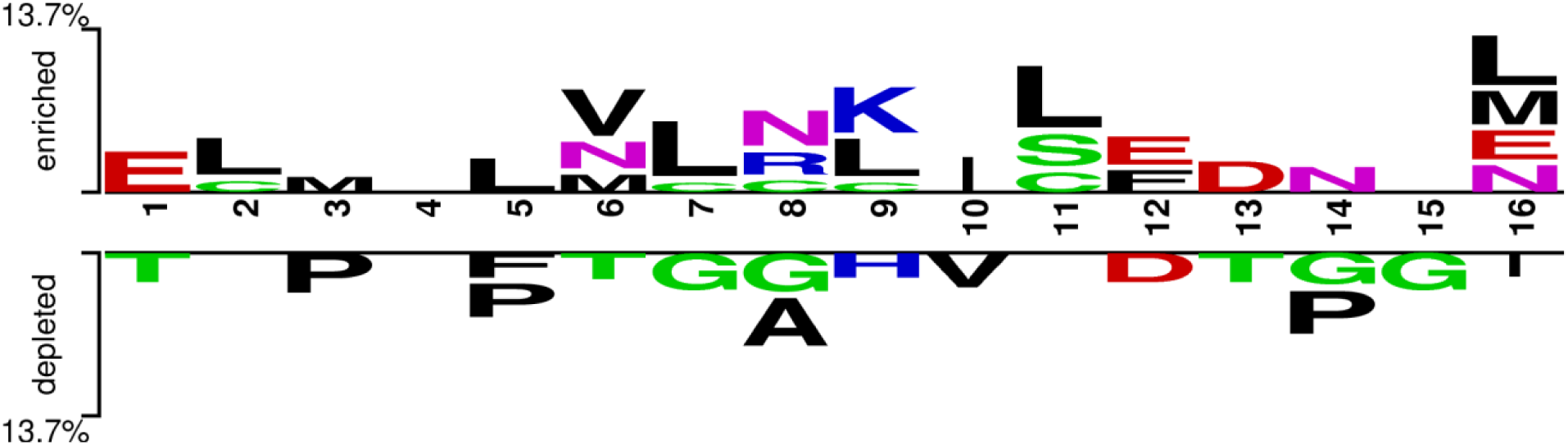
Two-sample logo displaying the amino acid positional conservation among IL-13 positive and negative dataset.

### Machine Learning Models Performance evaluation

In order to develop prediction models, we have used a positive dataset of 313 IL-13 inducing and a negative dataset of 2908 non-inducing unique peptides. We have applied various machine learning techniques such as SVM, KNN, LR, DT, RF, XGB, and GNB. The performance of all the developed models for top 95 features are reported in Table 2. RF based model outperforms other models with the AUC of 0.87 and 0.83 on the training and testing dataset, respectively.

**Table 2:**
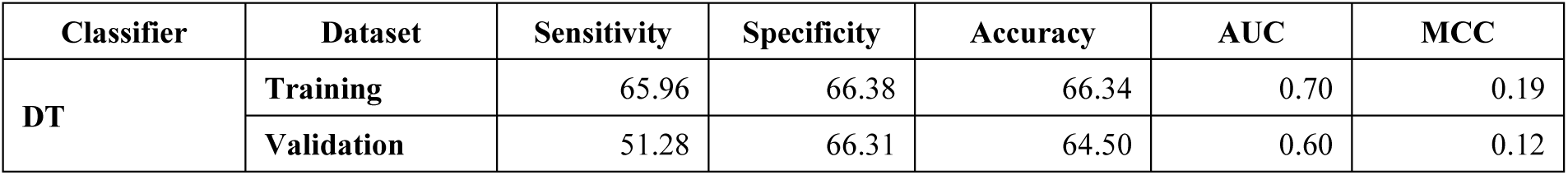
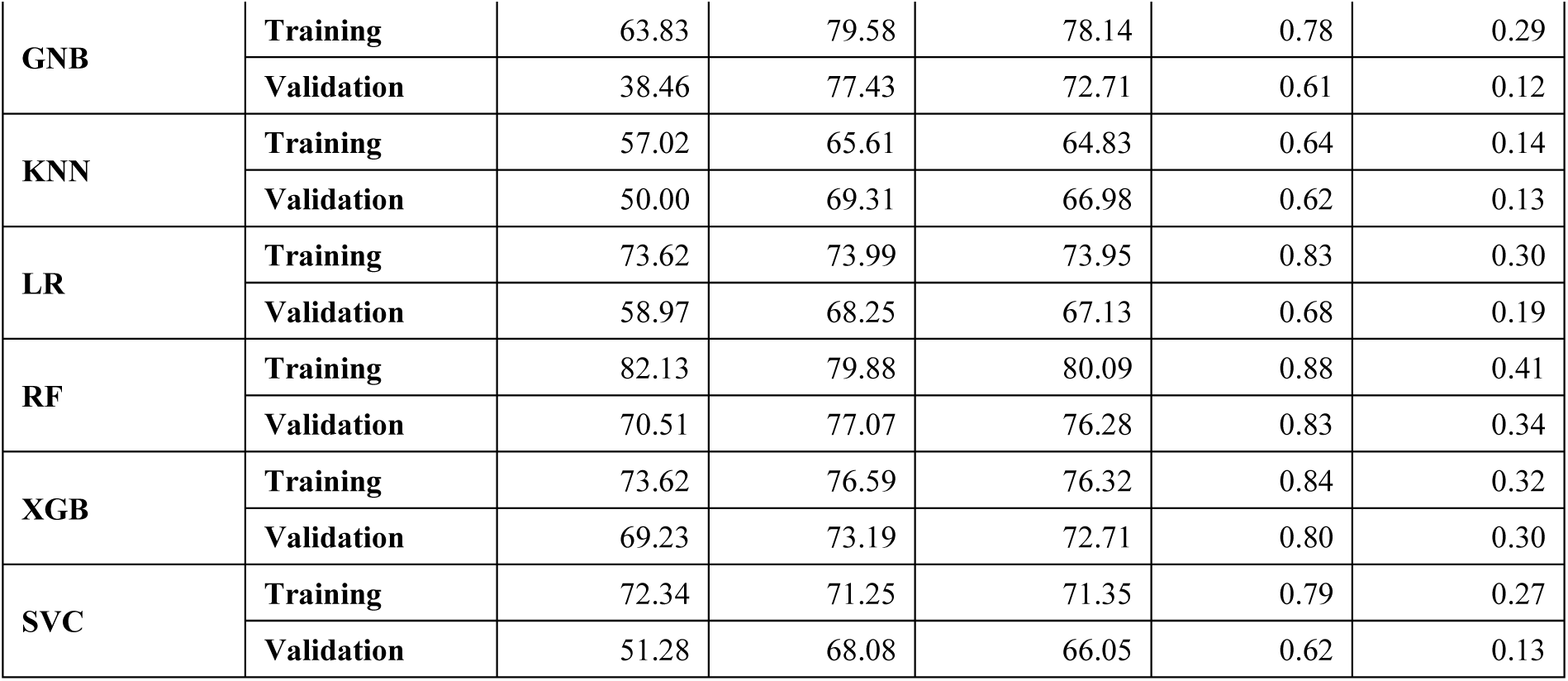
The performance of the machine learning models developed using top 95 features.

Further, we ranked the 95 selected features using feature-selector library of Python. In order to identify the best set of features without compromising the performance, we have developed the models on top 10, 20, 30, …95 features (Supplementary Table S2, S3). It was observed that the models developed on top-10 features performed quite well without much compromising the performance, as compared to the models developed on 95 features. As shown in Table 3, XGB based model outperforms other classifiers with AUC value of 0.83 and 0.80 on the training and testing dataset, respectively.

**Table 3:** The performance of the machine learning models developed using top 10 features.

| **Classifier** | **Dataset** | **Sensitivity** | **Specificity** | **Accuracy** | **AUC** | **MCC** |
| --- | --- | --- | --- | --- | --- | --- |
| **DT** | **Training** | 69.36 | 69.46 | 69.45 | 0.74 | 0.24 |
|  | **Validation** | 60.26 | 71.43 | 70.08 | 0.72 | 0.22 |
| **GNB** | **Training** | 71.06 | 68.18 | 68.44 | 0.74 | 0.24 |
|  | **Validation** | 64.10 | 66.31 | 66.05 | 0.73 | 0.21 |
| **KNN** | **Training** | 65.11 | 56.47 | 57.26 | 0.64 | 0.13 |
|  | **Validation** | 60.26 | 55.73 | 56.28 | 0.61 | 0.11 |
| **LR** | **Training** | 64.26 | 61.85 | 62.07 | 0.67 | 0.15 |
|  | **Validation** | 56.41 | 59.61 | 59.23 | 0.63 | 0.11 |
| **RF** | **Training** | 74.47 | 76.34 | 76.17 | 0.83 | 0.33 |
|  | **Validation** | 64.10 | 75.13 | 73.80 | 0.77 | 0.28 |
| **XGB** | **Training** | 78.30 | 74.84 | 75.16 | 0.83 | 0.33 |
|  | **Validation** | 71.80 | 73.02 | 72.87 | 0.80 | 0.30 |
| **SVC** | **Training** | 54.47 | 70.70 | 69.22 | 0.67 | 0.16 |
|  | **Validation** | 46.15 | 74.25 | 70.85 | 0.64 | 0.15 |

In order to check the robustness of the developed models, we have reshuffled the dataset 10 times and calculated the performance of models on top-10 features. Table 4 represents the mean and standard deviation of performance obtained after reshuffling the data ten times. We have observed that the performance of the models is maintained even after reshuffling, signifying the robustness of the developed models. XGB based model has shown the similar trend as in Table 3, with AUC of 0.82 ± 0.03

**Table 4:** The performance of the machine learning models developed after reshuffling dataset 10 times using top 10 features.

| Classifier | Sensitivity | Specificity | Accuracy | AUC | MCC |
| --- | --- | --- | --- | --- | --- |
| **DT** | $63.65 \pm 4.76$ | $67.94 \pm 2.83$ | $67.52 \pm 2.52$ | $0.71 \pm 0.03$ | $0.20 \pm 0.03$ |
| **GNB** | $72.38 \pm 4.92$ | $63.81 \pm 2.52$ | $64.65 \pm 2.30$ | $0.74 \pm 0.02$ | $0.22 \pm 0.03$ |
| **KNN** | $64.44 \pm 5.35$ | $51.25 \pm 2.00$ | $52.54 \pm 1.83$ | $0.62 \pm 0.04$ | $0.09 \pm 0.03$ |
| **LR** | $62.22 \pm 4.66$ | $60.76 \pm 1.87$ | $60.90 \pm 1.62$ | $0.66 \pm 0.02$ | $0.14 \pm 0.03$ |
| **RF** | $74.13 \pm 6.17$ | $73.95 \pm 1.41$ | $73.97 \pm 1.08$ | $0.81 \pm 0.03$ | $0.30 \pm 0.03$ |
| **XGB** | $76.98 \pm 4.98$ | $71.13 \pm 1.75$ | $71.71 \pm 1.51$ | $0.82 \pm 0.03$ | $0.31 \pm 0.03$ |
| **SVC** | $63.81 \pm 8.46$ | $63.20 \pm 3.35$ | $63.26 \pm 2.70$ | $0.68 \pm 0.04$ | $0.16 \pm 0.04$ |

### Case Study 1: IL-13 inducing peptides in proteins of SARS-CoV-2

Recent studies have reported the role of elevated IL-13 levels in COVID-19 patients [26–28]. Coronavirus proteins massively induces the release of this immunoregulatory cytokine. To identify the IL-13 inducing peptides in the SARS coronavirus proteins, we have utilized “Protein Scan” module of ‘IL-13Pred’ (https://webs.iiitd.edu.in/raghava/il13pred/scan.php) using the default parameters. We downloaded the SARS-CoV-2 proteins of five different countries, including India (MZ340539.1), China (MT291829.1), USA (MZ319836.1) Brazil (MZ169911.1) and Japan (LC528233.2) from NCBI (https://www.ncbi.nlm.nih.gov/datasets/coronavirus/). We identified 213 IL-13 inducing peptides out of 1259 peptides of the spike proteins of USA (Geolocation: California) reported SARS-CoV-2 strain (as reported in Supplementary Table S4). In addition to these, we have also identified the IL-13 inducing/non-inducing peptides in other coronavirus proteins such as envelope protein, ORF6, ORF1ab, ORF3a, ORF7a/7b, ORF8, nucleocapsid phosphoprotein, ORF10 and membrane glycoprotein of USA strain, represented in Supplementary Table S5. These findings shall be taken into consideration by the scientific community, while designing subunit vaccine against COVID-19 and other disorders that can be provoked by the induction of IL-13.

### Case Study 2: IL-13 inducing peptides in Spike proteins of SARS-CoV-2 variant strains

In this study, we have made a systematic attempt to highlight contrast among IL-13 inducing peptides in SARS-CoV-2 variant strains. We have identified and reported IL-13 inducing peptides in Alpha (B.1.1.7), Beta (B.1.351) and Delta (B.1.617.2) variants of SARS-CoV-2 virus. We have downloaded the reference Spike protein of SARS-CoV-2 from NCBI. Post that we have manually done the substitution mutations reported by Centers for Disease Control and Prevention (CDC portal) and Indian SARS-CoV-2 Genomics Consortium using In-House script. The Alpha variant (B.1.1.7) has total of seven substitution mutations namely N501Y, A570D, D614G, P681H, T716I, S982A and D1118H. The Beta variant (B.1.351) was obtained by doing subsequent nine substitution mutation D80A, D215G, K417N, E484K, N501Y, D614G, A701V, L18F and R246I in reference spike protein sequence. The Delta (B.1.617.2) variant of SARS-CoV-2 strain had nine mutations namely T19R, T95I, G142D, R158G, L452R, T478K, D614G, P681R and D950N from reference protein. In order to identify the IL-13 inducing peptides in these curated variants of COVID-19 Spike proteins, we have executed “Protein Scan” module of ‘IL-13Pred’ (https://webs.iiitd.edu.in/raghava/il13pred/scan.php) using the default parameters. Initially, we have computed IL-13 inducing and non-inducing peptides of all three variants strains. Subsequently, we have mapped the differences of three variants across the reference protein sequence in terms of IL-13 inducing vs non inducing properties. Table 6 represents the sequences with contrasting effects after substitution mutation in spike proteins. As shown in table below, few mutations have turned on the IL-13 inducing property and some have turned off the same. For instance, mutation at 618^th^ position of the spike protein in Delta strain from amino acid Proline to Arginine have turned on the IL-13 inducing property of the mutated peptide. Similarly, mutation from Aspartic acid to Asparagine at 950^th^ position have changed the nature of peptides from non-inducer to IL-13 inducer.

**Table 5:**
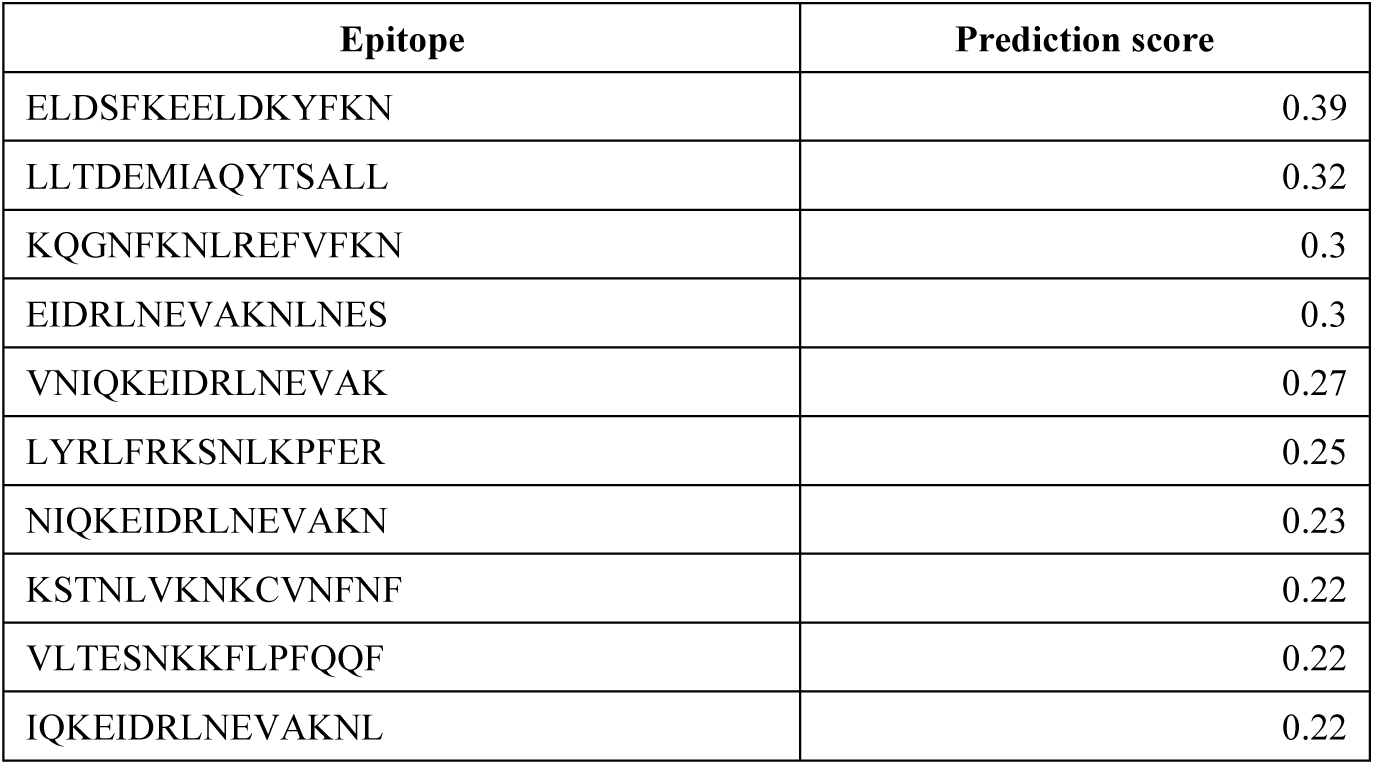
Potential IL-13 inducing peptides predicted by IL-13Pred in spike protein of USA (MZ319836.1) strain

**Table 6:**
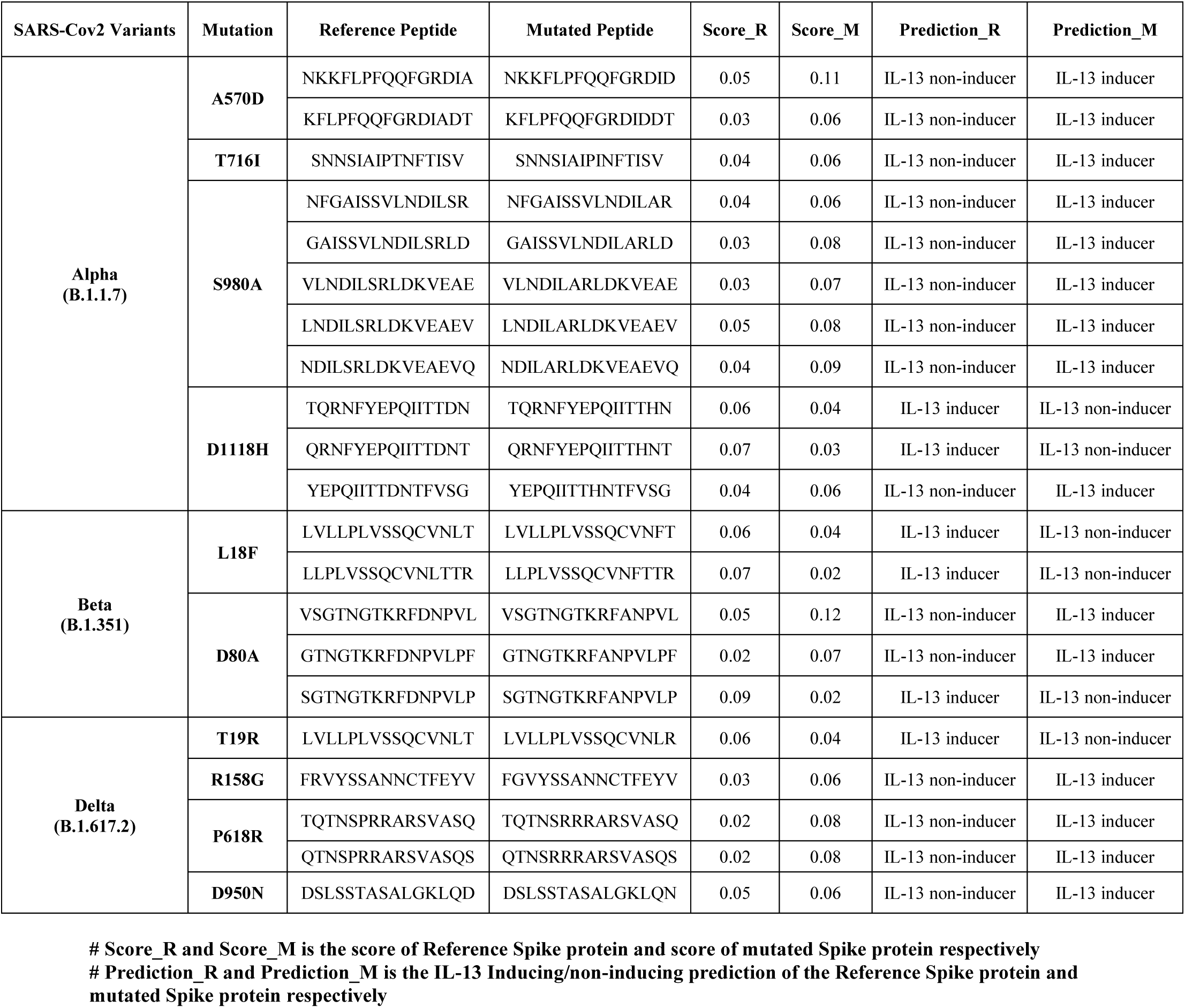
Changes in IL-13 inducing properties by mutation in Spike protein in SARS-CoV-2 variant strains.

However, the similar trends of these amino acid is depicted in Figure 3, where the composition of Arginine and Asparagine is preferred in IL-13 inducing peptides as compared to non-inducing peptides. In a given sequence, composition and position of the amino acid plays a vital role in changing the properties and role of the proteins in variant strains, which is also depicted in Table 6. The complete results of three variant spike protein strains are provided in the Supplementary Tables (S6-S8).

## Discussion and Conclusion

IL-13 have been shown its role in mucosal inflammation, including allergic asthma, eosinophilic oesophagitis, ulcerative colitis, and fibrosis [4]. We have also seen that receptors of IL-13 were found to be overexpressed in various cancer cells, defining it as a promising target for cancer immunotherapy. In recent study, researcher have shown the correlation between IL-13 levels and COVID-19 severity. Higher level of IL-13 has been reported in severe COVID-19 patients. They have also observed that COVID-19 patients with higher IL-13 serum levels tend to prolong ventilation support as compared to others. They have shown that IL-13 neutralization results in reduced COVID-19 severity and lung hyaluronan deposition in mouse model. In addition, they have reported that COVID-19 patients taking Dupilumab medicine had less severe disease [26]. In present study, we made a systematic approach to understand the role of IL-13 inducing peptides and have provided a method to classify IL-13 inducing peptides. The experimentally validated IL-13 inducing peptides of humans from IEDB are taken for developing prediction models. The positive dataset of 313 IL-13 inducing and a negative dataset of 2908 IL-13 non-inducing unique peptides, were used in this study. The amino acid composition and positional analysis was performed to show contrast among IL-13 inducing and non-inducing peptides. From compositional analysis we observe that the amino acid leucine, lysine, methionine and asparagine residues are preferred in IL-13 inducing peptides. However, glycine, proline, tyrosine and threonine residues are abundantly found in non-inducing peptides. On the other hand, the positional analysis shows that leucine is preferred at 2^nd^, 5^th^, 7^th^,9^th^, 11^th^ and 16^th^ position, whereas, asparagine is mostly preferred at 6^th^, 8^th^, 14^th^ and 16^th^ position in the positive dataset.

Further, we have applied Pfeature tool for extracting relevant features from this dataset. Using the extracted feature set, we have applied well-established machine learning methods with tuned parameters to develop the models with best performance. Machine learning model performance for both set of features, such as 95 selected features and top 10 features are reported. XGBoost-based model achieves a maximum AUC of 0.83 and 0.80 on training and independent dataset, respectively using top10 features. We have provided a webserver and a standalone version of ‘IL-13Pred’ for scientific community usage for distinguishing the IL-13 inducing and non-inducing peptides/epitopes. Using the protein scan module of ‘IL-13Pred’ webserver, we identify the IL-13 inducing peptides in 5 different COVID-19 strains such as India (MZ340539.1), China (MT291829.1), USA (MZ319836.1) Brazil (MZ169911.1) and Japan (LC528233.2). Also, we have identified the IL-13 inducing regions in Spike protein of Alpha (B.1.1.7), Beta (B.1.351) and Delta (B.1.617.2) variants of SARS-CoV-2 virus. The prediction results show that the amino acid P168R and D950N mutations in Delta variant induce IL-13 immunoregulatory cytokine.

The efficacy of an epitope/peptide to induce IL-13 and alter the immune response towards a disease state makes it of great importance in immunotherapy and vaccine designing. Although inducing IL-13 response in patients is a very complex process that depends on various factors. An epitope/ peptide is still a promising alternative while designing a vaccine or immunotherapy against any disease. In literature, studies have shown that antibodies that can block IL-13 receptors could be used in designing vaccine effectively [36,37]. Thus, IL-13 induced immunosuppression could serve as a crucial step in vaccine subunit designing. Although diverse *in silico* methods are available for T cell epitopes prediction, but no computational method was there for IL-13 inducing epitopes/ peptide prediction. The present study is an organized attempt made in this direction for providing a user-friendly platform/ webserver. We encourage scientific community to use IL-13Pred for developing efficient immunotherapy and vaccine candidates by differentiating them apriori as IL-13 inducing epitopes.

## Conflict of Interest

The authors declare no competing financial and non-financial interests.

## Author Contributions

SJ, AD, and SP collected and processed the datasets. SJ, AD, SP and GPSR implemented the algorithms. SJ, AD, and SP developed the prediction models. SJ, AD, SP and GPSR analysed the results. SP, SJ and AD created the back-end of the web server and front-end user interface. SJ, AD, SJ and GPSR penned the manuscript. GPSR conceived and coordinated the project, and gave overall supervision to the project. All authors have read and approved the final manuscript.

## Acknowledgement

AD is thankful to the Department of Science and Technology (DST-INSPIRE) and SP is thankful to the Department of Biotechnology (DBT) for providing senior research fellowships. SJ, AD and SP are thankful to the Department of Computational Biology, IIITD New Delhi for infrastructure and facilities.

## Data Availability Statement

All the datasets generated for this study are either included in this article the dataset is available at https://webs.iiitd.edu.in/raghava/il13pred/dataset.php.

## Notes

### Competing Interest Statement

The authors have declared no competing interest.

